# Cisplatin Induces BDNF Downregulation in Middle-Aged Female Rat Model while BDNF Enhancement Attenuates Cisplatin Neurotoxicity

**DOI:** 10.1101/2023.05.15.540850

**Authors:** Naomi Lomeli, Diana C. Pearre, Maureen Cruz, Kaijun Di, Daniela A. Bota

## Abstract

Cancer-related cognitive impairments (CRCI) are debilitating consequences of cancer treatment with platinum agents (e.g., cisplatin) that greatly alter cancer survivors’ health-related quality of life. Brain-derived neurotrophic factor (BDNF) plays an essential role in neurogenesis, learning, and memory, and the reduction of BDNF is associated with the development of cognitive impairment in various neurological disorders, including CRCI. Our previous CRCI rodent studies have shown that cisplatin reduces hippocampal neurogenesis and BDNF expression and increases hippocampal apoptosis, which is associated with cognitive impairments. Few studies have reported on the effects of chemotherapy and medical stress on serum BDNF levels and cognition in middle-aged female rat models. The present study aimed to compare the effects of medical stress and cisplatin on serum BDNF levels and cognitive performance in 9-month-old female Sprague Dawley rats to age-matched controls. Serum BDNF levels were collected longitudinally during cisplatin treatment, and cognitive function was assessed by novel object recognition (NOR) 14 weeks post-cisplatin initiation. Terminal BDNF levels were collected ten weeks after cisplatin completion. We also screened three BDNF-augmenting compounds, riluzole, ampakine CX546, and CX1739, for their neuroprotective effects on hippocampal neurons, *in vitro*. We assessed dendritic arborization by Sholl analysis and dendritic spine density by quantifying postsynaptic density-95 (PSD95) puncta. Cisplatin and exposure to medical stress reduced serum BDNF levels and impaired object discrimination in NOR compared to age-matched controls. Pharmacological BDNF augmentation protected neurons against cisplatin-induced reductions in dendritic branching and PSD95. Ampakines (CX546 and CX1739) but not riluzole altered the antitumor efficacy of cisplatin in two human ovarian cancer cell lines, OVCAR8 and SKOV3.ip1, *in vitro.* In conclusion, we established the first middle-aged rat model of cisplatin-induced CRCI, assessing the contribution of medical stress and longitudinal changes in BDNF levels with cognitive function. We conducted an *in vitro* screening of BDNF-enhancing agents to evaluate their potential neuroprotective effects against cisplatin-induced neurotoxicity and their effect on ovarian cancer cell viability.

## 2. Introduction

The first aim of this paper is to describe an aged female rat model of cisplatin-induced CRCI and examine the influence of medical stress and cisplatin on serum BDNF levels and cognitive function. The second aim is to conduct an in vitro screening of BDNF-enhancing pharmacological agents in primary rat hippocampal neurons to assess their neuroprotective effects against cisplatin-induced morphological damage. Lastly, the third aim is to screen these compounds in human ovarian cancer cell lines to assess their effect on cisplatin’s anti-cancer efficacy.

## 3. Materials and Methods

### 3.1 Animals

Animal studies were performed in accordance with the guidelines established by the NIH and the Institutional Animal Care and Use Committee (IACUC) of the University of California, Irvine. All animal experiments were approved by IACUC. All the data were generated using Sprague Dawley rats (Charles River Laboratories). Brain-derived neurotrophic levels have been negatively associated with cognitive function in cancer patients(1). Low BDNF levels have been associated with CRCI (cancer-related cognitive impairment) in female breast cancer survivors (2, 3). Breast and ovarian cancer are most commonly diagnosed in women peri-menopausal to menopausal aged 50-69 years. The median age at diagnosis for female breast cancer is 62 years(4), and 63 years for ovarian cancer(5).

In this study, twenty-eight, nine-month-old female retired breeder Sprague Dawley rats weighing 409 ± 45 grams at the start of the study served as subjects. Rats were group housed (4 rats per cage) or housed in pairs, kept on a standard light-dark cycle (12 h each) at 23°C ± 1°C room temperature, and 45% ± 10% humidity and given a standard rodent chow diet (Envigo Teklad 2920X) by the University Laboratory Animal Resources (ULAR).

Rats were randomized into three groups: Control (n=8), Saline (n=8), and Cisplatin (n=12). Rats were injected intraperitoneally with 2.5 mg/kg CDDP (Fresenius Kabi, USA, LLC.) dissolved in 0.9% saline every 2 weeks, for a total of 7 cycles, and a cumulative dose of 17.5 mg/kg CDDP. The vehicle animals received 0.9% sterile saline, i.p. of the same volume. Mannitol (APP Pharmaceuticals, 250 mg/kg, i.p.) was administered to all SAL and CDDP animals 1 h prior to CDDP administration to minimize renal toxicity and increase diuresis. Control rats did not receive any injections and were not subject to the medical stress protocol.

### 3.2 Medical Stress Protocol

Cancer incidence increases with age, and stress related to cancer diagnosis and treatment is common and negatively impacts the mental health and quality of life of patients(6). To examine the effects of cisplatin chemotherapy and medical stress on serum BDNF levels, anxiety, and cognitive function in 9-month-old Sprague Dawley retired breeders, we developed a medical stress protocol to model the effects of environmental stress associated with cancer care. Rats were randomized into three groups, healthy controls (CON), Saline (SAL), and Cisplatin (CDDP). The rats were allowed to acclimate to the vivarium for two weeks. CDDP and SAL rats received 5 ml of 0.9% saline, s.q., 3 times a week during each CDDP cycle to reduce dehydration associated with CDDP. Three days post-CDDP administration, blood draws from the lateral tail vein were collected to assess serum BDNF levels in the CDDP and SAL rats. The rats were restrained using a Tailveiner® restrainer (Braintree Scientific). A maximum of two blood draws were attempted per rat per timepoint. Subjects were weighed three times a week during the study. CON rats served as healthy controls and were not exposed to the medical stress protocol.

### 3.3 Serum BDNF Measurements

To assess the effect of cisplatin on peripheral BDNF levels, blood was collected from the lateral tail vein of VEH and CDDP rats 72 h after each CDDP cycle. Approximately 50 - 150 µL blood was collected per rat. Blood collection was attempted twice per rat per timepoint in the CDDP and SAL groups. Serum BDNF quantification in Fig 1.H-J, represents timepoints at which at least 3 serum samples were successfully collected and measured per group. SAL and CDDP serum BDNF levels were compared to the serum BDNF levels of healthy age-matched controls (CON) collected at the terminal timepoint. Blood was clotted for 30 minutes at room temperature (25°C) in 0.6 mL tubes, then centrifuged at 2,000 g for 15 minutes at room temperature.

**Figure 1.**
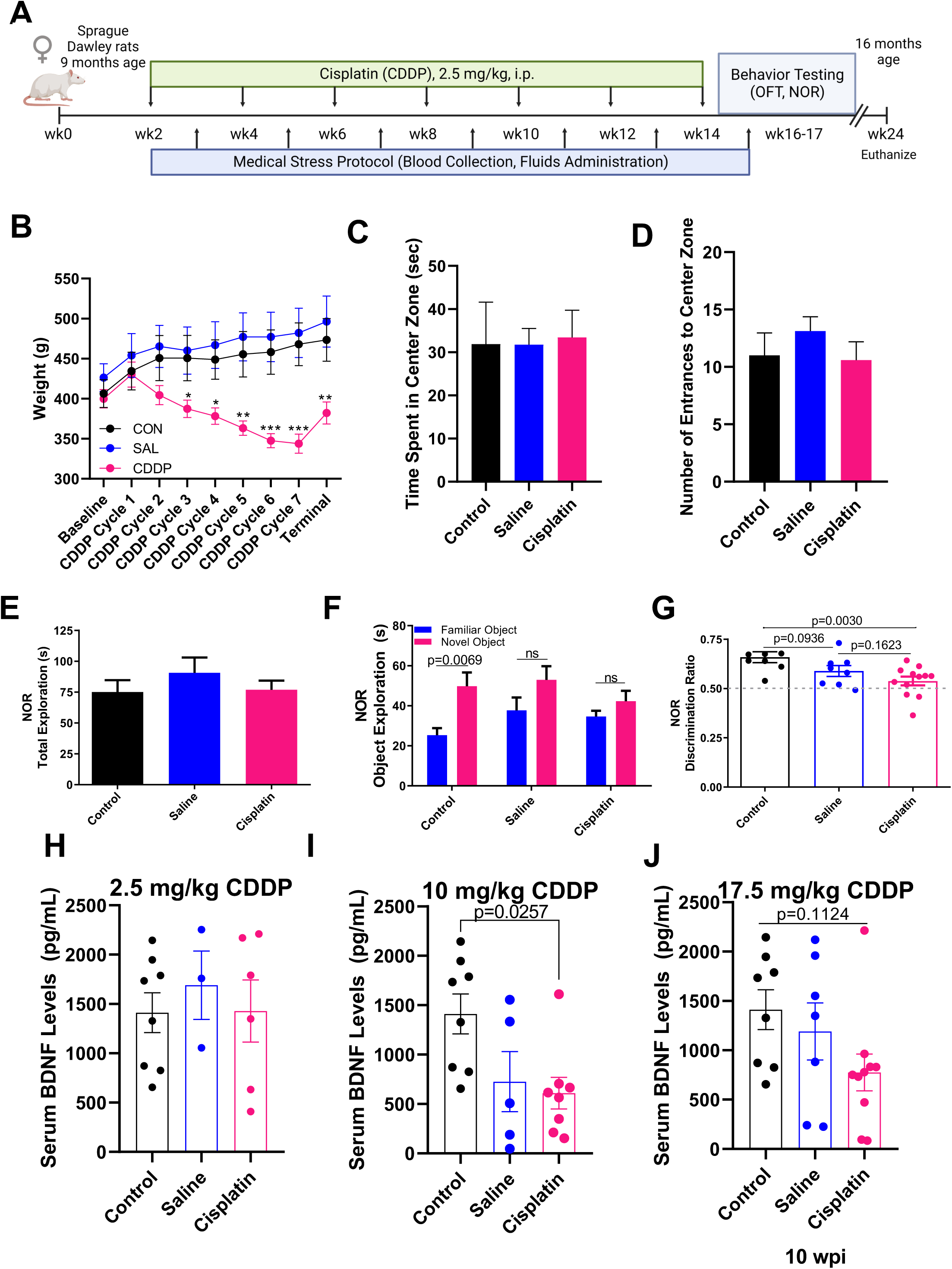
Effects of chronic CDDP exposure and medical stress induces on cognitive function and serum BDNF levels in female middle-aged rat model. (**A**) Schematic representation of the experimental design. 9- month-old female retired-breeder Sprague Dawley rats were injected with vehicle (SAL, saline, n=8) or cisplatin (CDDP, 2.5 mg/kg, i.p., n=12) once every two weeks, for 13 weeks. CDDP and SAL rats were subjected to the medical stress protocol, which included hydration therapy (5-10 mL 0.9% saline/day, s.q.) for five consecutive days following CDDP, and lateral tail vein blood collections 72 hours after each CDDP dose. Cage mates that did not receive CDDP/SAL injections nor blood collections served as healthy age-matched controls (CON, n=8) for this study. Fourteen weeks after the start of CDDP treatment, all rats were assessed for anxiogenic behavior using the open field recognition test (OFT), and cognitive function using the novel object recognition task (NOR). Rats were euthanized ten weeks following the completion of CDDP and serum was collected for BDNF analyses. (**B**) Effect of CDDP regimen on body weight (grams) over time. (**C**) Quantification of (**C**) time spent in the center zone and (**D**) number of entrances to the center zone of the arena over 10 minutes in the OFT. (**E**) Quantification of total time spent exploring novel and familiar objects during NOR. (**F**) Medical stress and CDDP treatment impaired cognitive function as SAL and CDDP rats had a reduced preference for the novel object comparted to the familiar object during NOR. (**G**) CDDP-exposed rats had a significantly lower discrimination ratio [(novel object exploration (sec))/(total object exploration (sec)] compared to healthy age-matched controls (CON). ELISA-based quantification of BDNF levels from rat serum showed longitudinal and dose-dependent changes in SAL and CDDP groups. Serum BDNF levels were quantified 72 hours following: (**H**) CDDP Cycle 1 (2.5 mg/kg, i.p.), (**I**) CDDP Cycle 4 (cumulative 10 mg/kg, i.p.) and (**J**) 10 weeks following CDDP Cycle 7 (cumulative 17.5 mg/kg, i.p.). Data are shown as mean ± SEM; each point represents one rat (**G-J**). ns, \**P*< 0.05, **P< 0.01, ***P<0.001, ****P<0.0001, as determined by one-way ANOVA with Tukey’s *post hoc* analysis for multiple comparisons test.

Serum (approximately 20-60 µl) was collected and stored at -80°C for further analysis. Serum concentrations of BDNF were measured using the Human/rat BDNF ELISA kit (Proteintech, KE00096) according to the manufacturer’s instructions at a 1:10 dilution. All samples from each subject were assayed simultaneously on the same plate and run in triplicate to minimize run-to-run variability.

### 3.4 Cognitive Testing

#### 3.4.1 Open Field Test

Open field testing was conducted 14 weeks after the initiation of cisplatin chemotherapy, and two weeks after cisplatin completion. To determine the effect of cisplatin on anxiogenic behavior after chronic chemotherapy, rats were placed in an open opaque square Plexiglas box arena (60 cm x 60 cm) with 60 cm high walls, at the center of the arena and allowed to freely explore for a 10 min session. Rats were removed from the arena at the end of the session, and the arena was cleaned with 70% ethanol, and dried before the subsequent session. Each session was recorded by an overhead video camera (Swann Security System) and analyzed manually by a blinded scorer. The arena was divided into two zones, a 30 cm x 30 cm center zone, and the periphery zone. Frequency of entrances and time spent in the center zone were quantified, with less time spent in the center zone considered anxiogenic behavior.

#### 3.4.2 Novel Object Recognition

To determine the effect of cisplatin on cognitive function after chronic chemotherapy, rats were examined on the novel object recognition (NOR) task 14 weeks after the initiation of cisplatin chemotherapy. We conducted NOR testing as previously described(7). Rats were placed in an open opaque Plexiglas box arena (60 cm x 60 cm) with 60 cm high walls containing two identical objects. Each rat explored the arena for 5 minutes per day for 2 consecutive days. The 5-minute test trial was given 24 h later, during which the rat was presented one of the familiar objects from the training phase paired with a new object. Total exploration time, time spent exploring each object (seconds), and the discrimination ratio (time spent exploring the novel object/total exploration time) were quantified.

### 3.5 Primary Rat Hippocampal Neuron Cultures

Hippocampal neuron cultures are prepared from postnatal day 0 (P0) Sprague Dawley pups as previously described(7-9). Cells were plated at a density of 7.5-10×10^3^ cells/coverslip on 12 mm coverslips (Chemglass Life Sciences) pre-coated with 0.2 mg/ml poly-D-lysine (Sigma Aldrich). Cells were maintained in Neurobasal Plus Medium (NBM+) with B-27 Plus (Gibco) at 37°C and 5% CO2. On day *in vitro* 3 (DIV3), cultures were treated with 1 µM arabinoside-cytosine (Sigma) to inhibit glial proliferation and refreshed twice a week with conditioned medium. Mature neurons were used for experiments on 17-21 days *in vitro* (17-21 DIV).

### 3.6 Ovarian Cancer Cell Lines

The established OVCAR8 and SKOV3.ip1 cell lines were maintained in RPMI 1640 medium with 300 mg/l L-Glutamine (Corning 10-040-CV) containing 10% FBS (Omega Scientific, Inc), and 1X penicillin/streptomycin (Gibco). These cell lines were kindly gifted by Dr. Olga Razorenova.

### 3.7 XTT Assay

The cells were seeded at approximately 1×104 cells/well in a final volume of 200 µl in clear 96-well plates. The plates were incubated at 37°C at 5% CO2 for 72 hr at the specified final concentrations of BDNF/CX546/Riluzole ± Cisplatin. Cell viability was determined using the Biotium XTT Cell Viability Kit (Biotium 30007).

### 3.8 Drug application in-vitro

*In-vitro,* recombinant human/murine/rat BDNF (Peprotech, 450-02) was reconstituted in 0.1% BSA in diH2O to make 100 µg/ml BDNF stocks, stored at -20°C. BDNF was diluted to a final concentration as specified in the respective cell culture medium. Riluzole (Sigma Aldrich, R116-25MG) was dissolved in DMSO to make a 10 mM working solution. Riluzole was freshly prepared immediately prior to each in vitro experiment, to minimize loss of efficacy after freeze-thawing. Ampakine CX546 (Tocris, 2980/10) was made into a 100 mM stock by dissolving in DMSO and stored at -20°C. Ampakine CX1739 was obtained from Cortex Pharmaceuticals. CX1739 was made into 25 mM stocks in 33% (2-Hydroxypropyl)-β-cyclodextrin/H_2_O/0.9% NaCl. For all experiments, all treatment groups were exposed to an equal volume of the respective vehicle, at the same timepoints.

### 3.9 Immunocytochemistry (ICC)

Neurons were fixed with ice-cold 4% paraformaldehyde (PFA, Thermo Scientific AC41678-00) in PBS, pH 7.4 for 12 min. To evaluate dendritic branching, neurons were immuostained with an antibody against postsynaptic density-95 (PSD95). The cells were incubated with mouse anti-PSD95 1:4000 (Thermo Fisher, MA1-046) in 200 µl per well in blocking buffer (0.3% FBS, 0.1% Triton-X in PBS, pH 7.4) at 4°C. The following day, coverslips were washed with PBS and incubated in goat anti-mouse Alexa Fluor 594 nm (Jackson Immuno Research, NC0540445) at room temperature for 1.5 h. Coverslips were mounted on slides using DAPI Fluoromount G (Southern Biotech) mounting medium.

### 3.10 PSD95 Puncta Quantification

Dendritic spine density was quantified as the number of PSD95 positive puncta on dendritic branches. Images were generated by confocal microscopy, Olympus FV3000. 3 µm z-series (0.5 µm steps) images were captured from dendrites that were clearly distinct form dendrites of other neurons and dendritic crossings at 60X 1.5 zoom (NA 1.42) using an oil-immersion objective. Each experiment included 3 coverslips per treatment group, with 4 dendrites from 2 separate neurons analyzed per coverslip, for a total of 6 dendrites per treatment group. Spine density was expressed as the number of PSD95 puncta per 20 µm of dendritic length, extending to 100 µm from the soma. Images were processed for analyses by conversion into 8-bit black and white TIFF files, then scaled for distance per pixel length using Fiji. The distance from the soma was then measured and divided into 20 µm segments. Each individual puncta were considered separate spines, the number of PSD95 puncta localized in clusters were adjusted based on the cluster size.

### 3.11 Sholl Analysis

Dendritic branching was evaluated using Sholl analysis. Total dendritic length was measured, and the number of intersections between branches and concentric circles at increasing 20 µm segments from the soma were quantified. Images were generated by confocal microscopy, Olympus FV3000. 8 µm z-series (1 µm steps) images were captured at 20X (NA 0.75) spanning across entire neurons. Each experiment included three coverslips per treatment group, with three neurons imaged per coverslip, for a total of nine images per treatment group. Images were processed for analyses by conversion into 8-bit black and white TIFF files, and concentric circles were overlayed on the images using the Concentric Circles plug-in in Fiji.

### 3.12 BDNF qPCR assay

Total RNA was extracted using RNeasy Mini Kit (Qiagen, Germantown, MD, USA), and cDNA was generated using the iScript cDNA Synthesis Kit (Bio-Rad, Hercules, CA, USA) as previously described(7). Quantitative PCR reactions (iQTM SYBR Green Supermix, Bio-Rad) were done using a Bio-Rad CFX96 Real-time System. *BDNF* gene expression levels were normalized to *GPADH* expression levels. The sequences for rat BDNF and GAPDH primer sets were: *BDNF CDS* forward 5′-AAACGTCCACGGACAAGGCA-3′, reverse 5′-TTCTGGTCCTCATCCAGCAGC-3′; *GAPDH* forward 5′-CCTTCATTGACCTCAACTACAT-3′, reverse 5′-CCAAAGTTGTCATGGATGACC-3′. The primers were ordered from IDT, Integrated Device Technology, Inc., Coralville, Iowa, USA.

### 3.13 Systematic analysis and statistical considerations

Each experiment included three replicates per treatment groups and was repeated twice. All imaging and analysis of PSD95 and dendritic branching, and behavioral testing were performed by two researchers blinded to the experimental conditions. Analyses of all biochemical, behavioral, and imaging experiments were performed using one-way ANOVA, or two-way repeated (RM)-ANOVA, followed by Tukey’s post-hoc multiple comparisons test. All data are presented as the mean±SEM and significance levels were set at 0.05. Data were analyzed using GraphPad Prism 8.0 Software.

## 4. Results

### Cisplatin and Medical Stress Decreases Serum BDNF and Impairs Cognitive Function in Female Rats

9-month-old female rats received chronic CDDP treatment (2.5 mg/kg, i.p.) every other week for 13 weeks, and received a cumulative CDDP dose of 17.5 mg/kg, i.p. (**Fig. 1A**). This CDDP regimen was well-tolerated with no adverse toxicities (i.e., weight loss >20%, death) (**Fig. 1B, Suppl. Fig. S1A**). Blood samples were collected 72 hours following CDDP injections in the SAL and CDDP groups. In addition, SAL and CDDP rats received 5-10 ml of 0.9% saline, s.q., for 5 days as hydration therapy following CDDP injections. This medical stress protocol models the clinical conditions cancer patients are exposed to during cancer treatment(10).

To determine the tolerable clinically-relevant CDDP regimen in this model that induces CRCI without adverse toxicities (i.e., weight loss >20%, death) we first conducted a pilot study using a chronic CDDP regimen of 5 mg/kg, i.p., every other week (n=12). The first dose resulted in a 14.3% decline in baseline weight, and significant toxicity was observed after the second cycle, therefore we dose-reduced and dose-delayed to allow weights to stabilized, and we determine that 2.5 mg/kg could be administered biweekly with minimal toxicity (**Suppl. Fig. S1B,C**). This regimen used in this study (2.5 mg/kg/biweekly) for 13 weeks is lower than that which induced CRCI in 2-month-old male Sprague Dawley rats (5 mg/kg, i.p.) weekly for 4 weeks(7). CDDP induced an additive reduction in body weight over time (*F*(16, 198) =15.03, *P* <0.0001, **Fig. 1B**) Renal toxicity is the one of the major dose-limiting side effects of CDDP, and compared to younger patients, the incidence of cisplatin-induced nephrotoxicity is increased in older patients, which may influence clinician’s decision to use a low-dose regimen of CDDP(11)>. Our objective was to assess the effects of medical stress in serum BDNF levels, anxiogenic behavior, and novel object recognition on rats receiving a CDDP regimen or saline, compared to healthy controls.

Two weeks after CDDP treatment, anxiogenic behavior was assessed using the open field test (OFT), followed by the novel object recognition (NOR) task to assess cognitive performance. Open field activity was observed on day 1 of the habituation phase. There was no significant difference in the time spent in the center zone of the arena (*F*(2, 25) = 0.01957, *P* = 0.9806) or the number of entrances to the center zone (*F*(2,25) = 0.6460, *P =* 0.5327) between the groups (**Fig. 1C,D**), which suggests the absence of anxiogenic behavior during the cognitive testing. During the NOR test phase, there was no difference in the total time spent exploring the novel and familiar object between the experimental groups (**Fig. 1E**). Comparison of the time spent exploring the novel object compared to the familiar object revealed significant differences in object exploration in the CON rats (*P =* 0.0069), but not for the SAL (*P* = 0.1256) or CDDP-treated rats (*P* = 0.2118, **Fig. 1F**). A one-way ANOVA revealed a significant overall treatment effect between groups (*F*(2,25) =5.897, *P*= 0.0080) . The discrimination ratio was calculated as [(time spent exploring the novel object)/(total exploration time)] for each subject. CDDP-treated rats showed a significantly reduced discrimination ratio compared to the CON rats who were not exposed to the medical stress protocol (*P* = 0.0030), but no difference between the SAL rats. SAL rats showed a trend towards reduced discrimination compared to the CON rats (*P =* 0.1623, **Fig. 1G**). These data suggest that CDDP results in cognitive deficits, and that medical stress associated with cancer treatment may influence these impairments.

### Effect of Chronic Cisplatin and Medical Stress in Serum BDNF Levels

Brain Derived Neurotrophic Factor (BDNF), a member of the neurotrophin family, plays a key role in promoting neuronal survival and growth, synaptogenesis, and is essential to learning and memory(12). Past studies in rat models have shown that cisplatin treatment reduces BDNF levels in the hippocampus(8, 13). Chronic stress is associated with loss of dendritic arborization and spines, changes in synaptic plasticity, and cognitive impairment(14). Although there are multiple factors involved in stress-induced changes in the plasticity and integrity of excitatory synapses(15), chronic stress induces transient downregulation of BDNF mRNA levels in the rat hippocampus(16). To examine the longitudinal effects of cisplatin and medical stress on serum BDNF levels and association with the behavioral outcomes observed in the SAL and CDDP-treated rats, we collected serum samples 72 hours after each CDDP cycle and 10 weeks post-CDDP completion, and measured BDNF levels by ELISA (**Fig. 1H-J**). One-way ANOVA revealed a significant overall treatment effect between groups (*F*(2,18) = 4.640, *P* = 0.0237) following CDDP Cycle 4 (**Fig. 1J**). Serum BDNF levels were significantly lower in CDDP-treated rats (609.2 pg/mL ± 161.1 pg/mL) compared to CON (1412 pg/mL ± 202 pg/mL), 72 hours following CDDP Cycle 4 (*P = 0.0257).* Serum BDNF levels were also decreased in SAL-treated rats (726.5 pg/mL ± 304.7 pg/mL) but not significantly compared to CON (*P = 0.1065)* (**Fig. 1J**). Notably, the serum BDNF levels partially recovered in SAL rats (1190 pg/mL ± 290.1 pg/mL), while the levels remained low in CDDP-treated rats (775.2 pg/mL ± 187 pg/mL) compared to CON (1412 pg/mL ± 202 pg/mL) (**Fig. 1H**). Although not significant these trends suggest that medical stress induces transient reductions in BDNF that may resolve following treatment completion, however the effects of CDDP on serum BDNF levels are dose-dependent and last long following treatment discontinuation.

### In vitro BDNF application prevents cisplatin-induced reductions in dendritic branching and PSD95 puncta density

Reductions in BDNF expression are associated with cognitive impairment in multiple models of CRCI (17-19). We previously reported that cisplatin significantly reduces BDNF mRNA levels. This reduction in BDNF expression was accompanied by reductions in postsynaptic density-95 (PSD95) density, a surrogate marker of dendritic spines, in rat hippocampal neurons following exposure to 0.1 µM, 1 µM at short (2 h) and chronic (24 h, 48 h) time-points (7, 8). To test directly whether BDNF supplementation could prevent cisplatin-induced reductions in dendritic branching and spine density, we assessed the effect of CDDP and BDNF administration on dendritic arborization via Sholl Analysis, and expression of PSD95 in mature rat cultured hippocampal neurons.

We measured the number of dendritic branch points from soma in hippocampal neurons exposed to 1 µM cisplatin for 24 h (**Fig. 2A, B**). Sholl analysis showed that cisplatin significantly reduced the number of dendritic intersections compared to vehicle (*P* = 0.0034), which was prevented by co-treatment with BDNF. Two-way repeated measures ANOVA revealed a significant interaction between cisplatin and BDNF, *F*(45,360) = 2.619, *P* < 0.0001. The cisplatin-induced loss of dendritic arborization was also accompanied by reduction in PSD95 density compared to vehicle (*P* <0.0012), which were prevented by BDNF (*F*(3,20) = 12.77, *P* < 0.0001, **Fig. 2C,D**).

**Figure 2.**
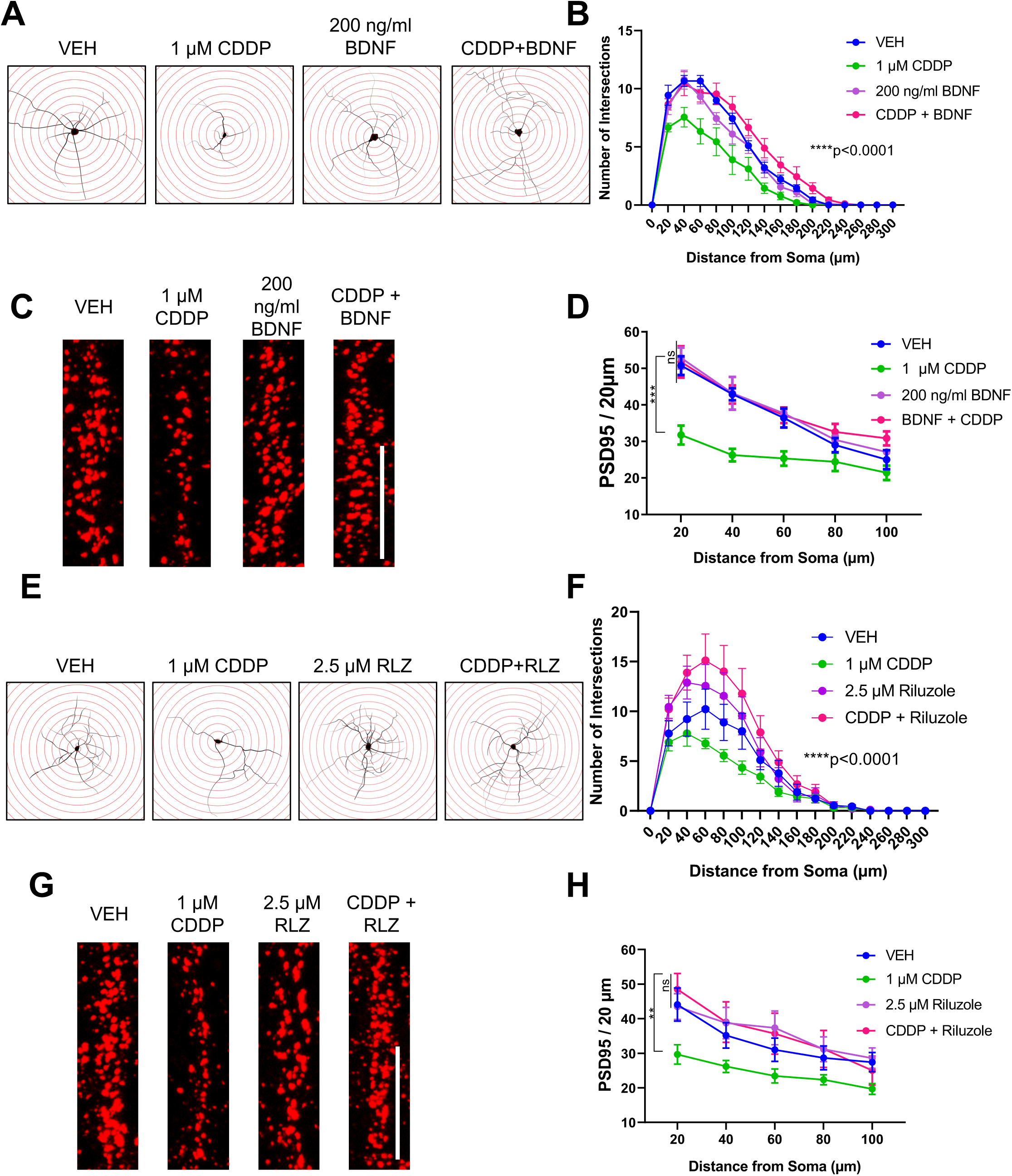
In vitro, BDNF augmentation is protective against cisplatin-induced loss of dendritic arborization and PSD95. (**A**) Mature rat hippocampal neurons were immuostained for PSD95 following exposure to 1 µM CDDP with or without co-treatment of 200 ng/mL BDNF for 24 h. Representative images of neurons reconstructed with Adobe Illustrator and superimposed over concentric Sholl circles (20 µm increments). (**B**) Quantification of intersection points between the concentric circles and dendritic branches at increasing distance from the soma. (**C**) Representative images and (**D**) quantification of PSD95 density at increasing distance from soma in neurons exposed to 1 µM CDDP with or without co-treatment of 200 ng/mL BDNF for 24 h. (**E**) Neurons were pre-treated with 2.5 µM RLZ for 1 h followed by exposure to 1 µM CDDP for 24 h. Representative images of reconstructed neurons superimposed over Sholl circles and (**F**) quantification of dendritic branch points using Sholl analysis. (**G**) Representative images and (**H**) quantification of PSD95 density in neurons pre-treated with 2.5, 5 µM RLZ for 1 h followed by exposure to 1 µM CDDP for 24 h. Data are shown as mean ± SEM; (**B,F**) n=9 neurons/group; (**D, H**) n= 6 neurons/group, 2 dendritic branches/neuron. Not significant = ns, \**P*< 0.05, \*\**P*< 0.01, \*\*\**P*<0.001, \*\*\*\**P*<0.0001, as determined by two-way repeated measures ANOVA with Tukey’s *post hoc* analysis for multiple comparisons test. Scale bars = 10 µm.

### In Vitro, BDNF prevents cisplatin-induced reductions in dendritic arborization and PSD95 density

Given the critical role of BDNF in regulating synaptic plasticity, and the association of low BDNF levels with the development of CRCI, we next sought to examine whether pharmacological enhancement of BDNF may be a novel therapeutic strategy to prevent CRCI. We conducted a screening study of three neuroprotective pharmacological agents that have been shown to increase BDNF and improve cognitive function: riluzole (RLZ), Ampakine CX546, and Ampakine CX1739. We first conducted a dose-response experiment to determine the effects of RLZ on dendritic arborization and PSD-95 density. We tested four doses 1 µM, 2.5 µM, 5 µM, and 10 µM RLZ (**Suppl. Fig. 2A,B**). Riluzole alone did not significantly change dendritic arborization (*F*(4,32) = 1.720, *P* = 0.1698) or PSD-95 density (*F*(4,25) = 0.6280, *P* = 0.6470) compared to vehicle at any of the tested doses at all distance points along the dendrites. In the presence of RLZ (1 hour pre-treatment), CDDP no longer reduced dendritic arborization (*F*(45,480) = 2.559, *P* <0.001, **Fig 2. E,F**). RLZ also protected dendritic spines from cisplatin-induced PSD-95 loss (*F*(12, 80) = 1.899, P = 0.0467, **Fig 2. G,H**).

### In Vitro, Ampakines CX546 and CX1739 protect against cisplatin-induced reductions in dendritic arborization and PSD95 density

Ampakines are positive allosteric modulators of AMPA receptors, and have been shown to up-regulate BDNF, rescue synaptic plasticity, and learning in rodent models of cognitive impairment(20-23). We had shown previously that cisplatin induced a pronounced reduction in BDNF mRNA levels in primary hippocampal neurons as early as 2 h after treatment(8). We next asked whether ampakine administration could prevent cisplatin-induced reductions in BDNF expression (**Fig. 3A**). 0.1 µM Cisplatin significantly reduced BDNF mRNA expression levels compared to VEH (*P* = 0.0032), 24 h post-treatment. Ampakine CX546 alone (50 µM) significantly increased BDNF levels by ∼6-fold compared to VEH (*P*<0.0001). Notably, CX546 significantly prevented cisplatin-induced reductions in BDNF levels compared to cisplatin alone (*P*<0.0001). One-way ANOVA revealed a significant treatment effect between cisplatin and CX546, *F*(3,8)= 407, *P*<0.0001).

**Figure 3.**
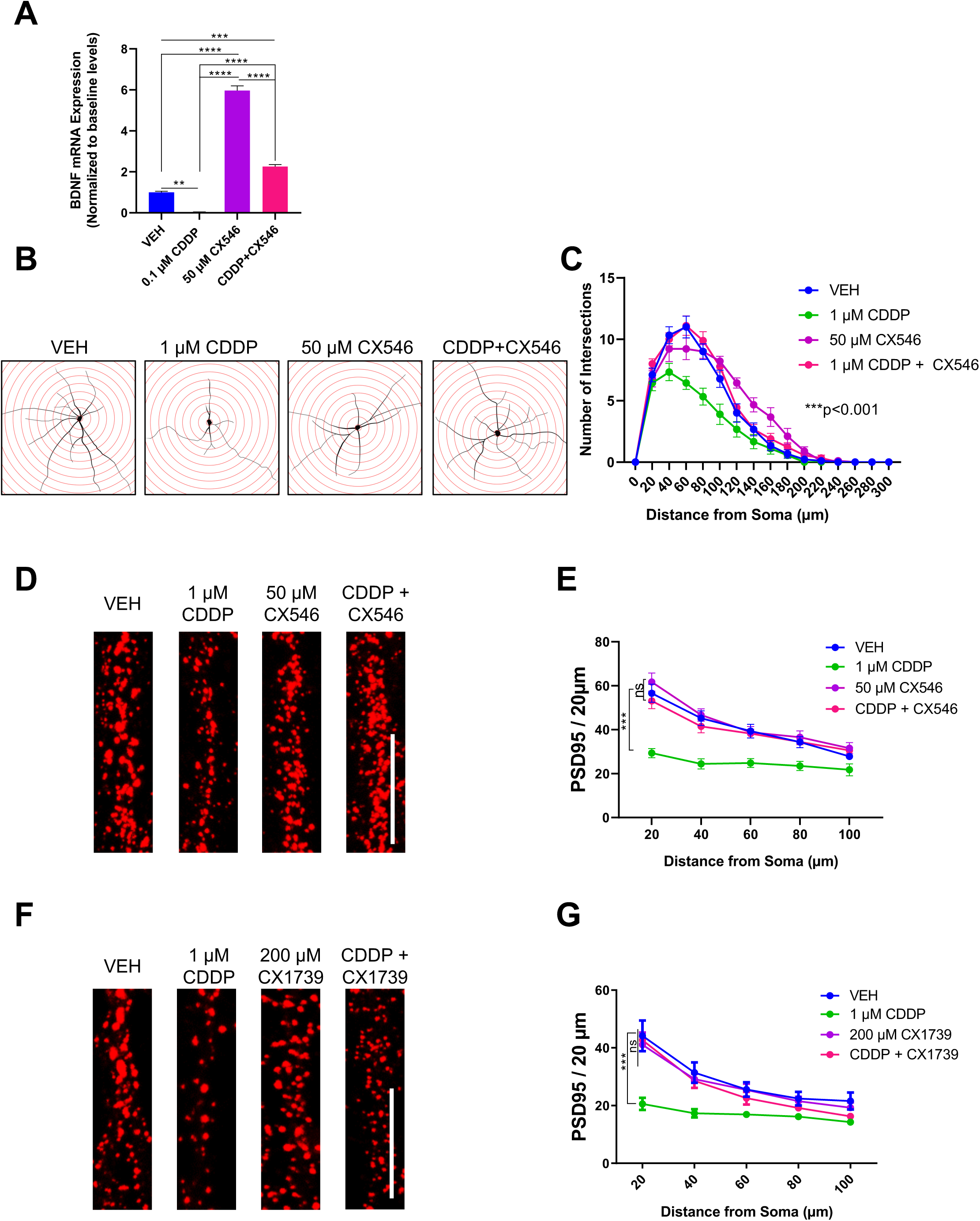
In vitro, ampakine CX546 prevents cisplatin-induced loss of BDNF expression, dendritic arborization, and PSD95. (**A**) Cisplatin significantly reduced BDNF mRNA after exposure to 1 µM cisplatin at 24 h, which was prevented by co-treatment with 50 µM CX546. (**B**) Representative images of reconstructed neurons superimposed over 20 µm concentric Sholl circles, and (**C**) Sholl analysis quantification of mature hippocampal neurons exposed to 1 µM cisplatin with or without co-treatment of 50 µM CX546 for 24 h. (**D**) Representative images of dendritic branches immuostained for PSD95 and (**E**) quantification of PSD95 puncta density along distance from soma following exposure to 1 µM CDDP with or without 50 µM CX546 for 24 h. (**F**) Representative images and (**G**) quantification of PSD95 puncta along distance from soma following 48 h co-treatment with 1 µM cisplatin and with or without 200 µM ampakine CX1739. (**C**) n=9 neurons/group; (**E,G**) n=6 neurons/group, 2 dendritic branches/neuron. Not significant = ns, \**P*< 0.05, \*\**P*<0.01, \*\*\**P*<0.001, \*\*\*\**P*<0.0001, as determined by two-way repeated measures ANOVA with Tukey’s *post hoc* analysis for multiple comparisons test. Scale bars = 10 µm.

Co-treatment with ampakine CX546 significantly prevented cisplatin-induced reductions in dendritic arborizations (*F*(45,360) = 4.570, *P*<0.0001, **Fig. 3B,C**). It also prevented cisplatin-induced reductions in PSD95 loss (*F*(12,80) = 2.864, *P* = 0.0025, **Fig. 3D,E**). Additional studies examining the effect of a different ampakine, CX1739 showed that CX1739 prevented cisplatin-induced PSD95 loss (*F*(12,80)=6.328, *P*<0.0001, **Fig. 3F,G**).

### In Vitro Screening of BDNF-enhancing compounds in human ovarian cancer cell lines

To examine whether BDNF alters cancer cell viability in vitro, we assessed viability of two human ovarian cancer cell lines OVCAR8 and SKOV3.ip1 exposed to 10 µM cisplatin with or without increasing concentrations of BDNF, Riluzole, CX546, or CX1739, for 72 h (**Fig. 4**). Cisplatin (10 µM) reduced OVCAR8 viability to 45.23% ± 0.9763 and 30.75% ± 0.5271% in SKOV3.ip1 at 72 h. BDNF alone had no effect on OVCAR8 and SKOV3.ip1 at BDNF concentrations ranging between 50 ng/ml to 200 ng/ml, with 300 ng/ml BDNF inducing a 4.84% ± 0.2447% reduction in SKOV3.ip1 viability (**Fig. 4A**). BDNF did not reduce cisplatin’s anti-cancer efficacy in cells co-treated with BDNF and cisplatin at any of the doses tested. Riluzole alone had no effect on cell viability compared to the vehicle control (**Fig. 4B**). The addition of riluzole (2.5 – 20 µM) to cisplatin-treated cancer cells did not affect cisplatin’s killing efficacy (**Fig. 4B**). At the highest concentration tested, 300 µM CX546, induced a significant increase in OVCAR8 (3.9% ± 0.3885%) and SKOV3.ip1 (7.6% ± 0.5028%) viability compared to the vehicle control (*P*<0.001, **Fig. 4C**). Co-treatment with CX546 reduced cisplatin’s anti-cancer efficacy (*P*<0.001). The addition of 50 µM CX 546 to cisplatin-treated OVCAR8 and SKOV3.ip1 increased cell viability to 10.79% and 10.88% compared to 10 µM cisplatin alone, respectively (*P*<0.001, *P*<0.001, **Fig. 4C**). This effect was dose dependent in the SKOV3.ip1 cell line as co-treatment with 300 µM CX546 dose exerted a 30.48% increase in viability compared to 10 µM cisplatin alone (*P*<0.001, **Fig. 4C**). Ampakine CX1739 exerted an increase in viability, with 300 µM CX1739 exerting a significant increase in OVCAR8 (24.2% ± 2.715%, p=0.0432) viability compared to the vehicle control (**Fig. 4D**). Unlike CX546, CX1739 did not alter cisplatin’s anti-cancer efficacy, when co-applied with cisplatin to ovarian cancer cells.

**Figure 4.**
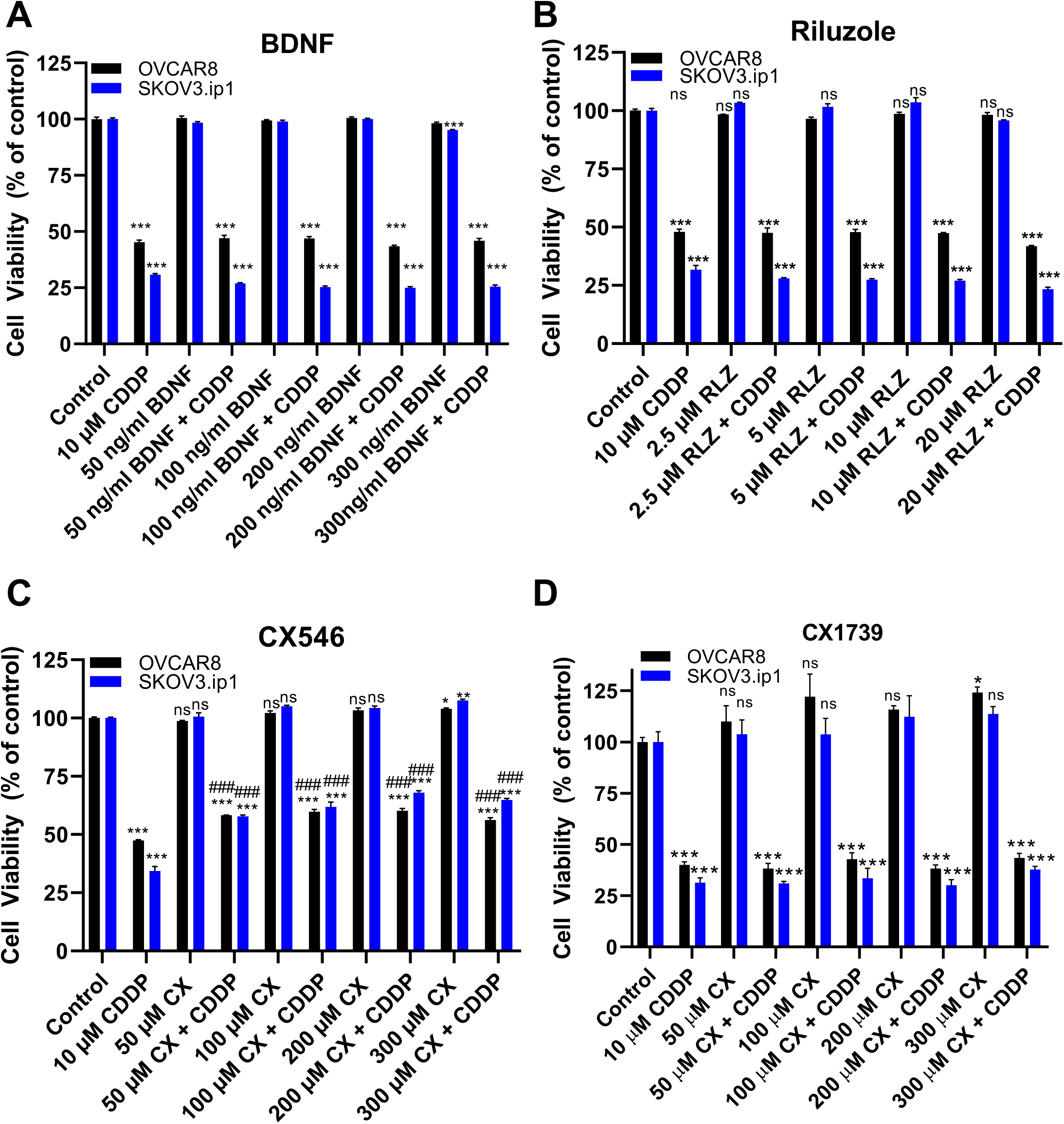
In vitro, screening of BDNF-enhancing pharmacological agents, riluzole, CX546, and CX1739 in human ovarian cancer lines OVCAR8 and SKOV3.ip. (**A**) OVCAR8 and SKOV3.ip1 were plated at 10,000 cells/well on 96-well plates, and treated with 10 µM cisplatin with or without graded doses of (A) BDNF (50 ng/ml, 100 ng/ml, 200 ng/ml, 300 ng/ml) (**B**) riluzole (2.5 µM, 5 µM, 10 µM, 20 µM), (**C**) CX546 (50 µM, 100 µM, 200 µM, 300 µM), and (**D**) CX1739 (50 µM, 100 µM, 200 µM, 300 µM) for 72 hours. Cell viability was normalized to vehicle control. (A-D) n=3 wells/group. Not significant = ns, \**P*<0.05, \*\**P*<0.01, *\*\*\*P*<0.001, \*\*\*\**P*<0.0001 denotes statistical significance compared to vehicle control. ^###^*P<0.001* denotes statistical significance compared to 10 µM cisplatin. Statistical significance was determined by one-way ANOVA with Tukey’s post hoc analysis for multiple comparisons test.

### Abbreviations

CDDP: Cisplatin;
RLZ: Riluzole;
PSD95: Postsynaptic density-95;
CRCI: Cancer-Related Cognitive Impairments;
BDNF: brain-derived neurotrophic factor;
SAL: saline;
VEH: vehicle;
NOR: novel object recognition;
OFT: open field test

## Acknowledgements

We thank Drs. Gary Lynch and Julie Lauterborn for providing CX1739 and advice with the ampakine experiments. We thank Dr. Senjie Du for his technical assistance with the BDNF experiments in hippocampal neurons.

---

**Supplementary Figure 1.**
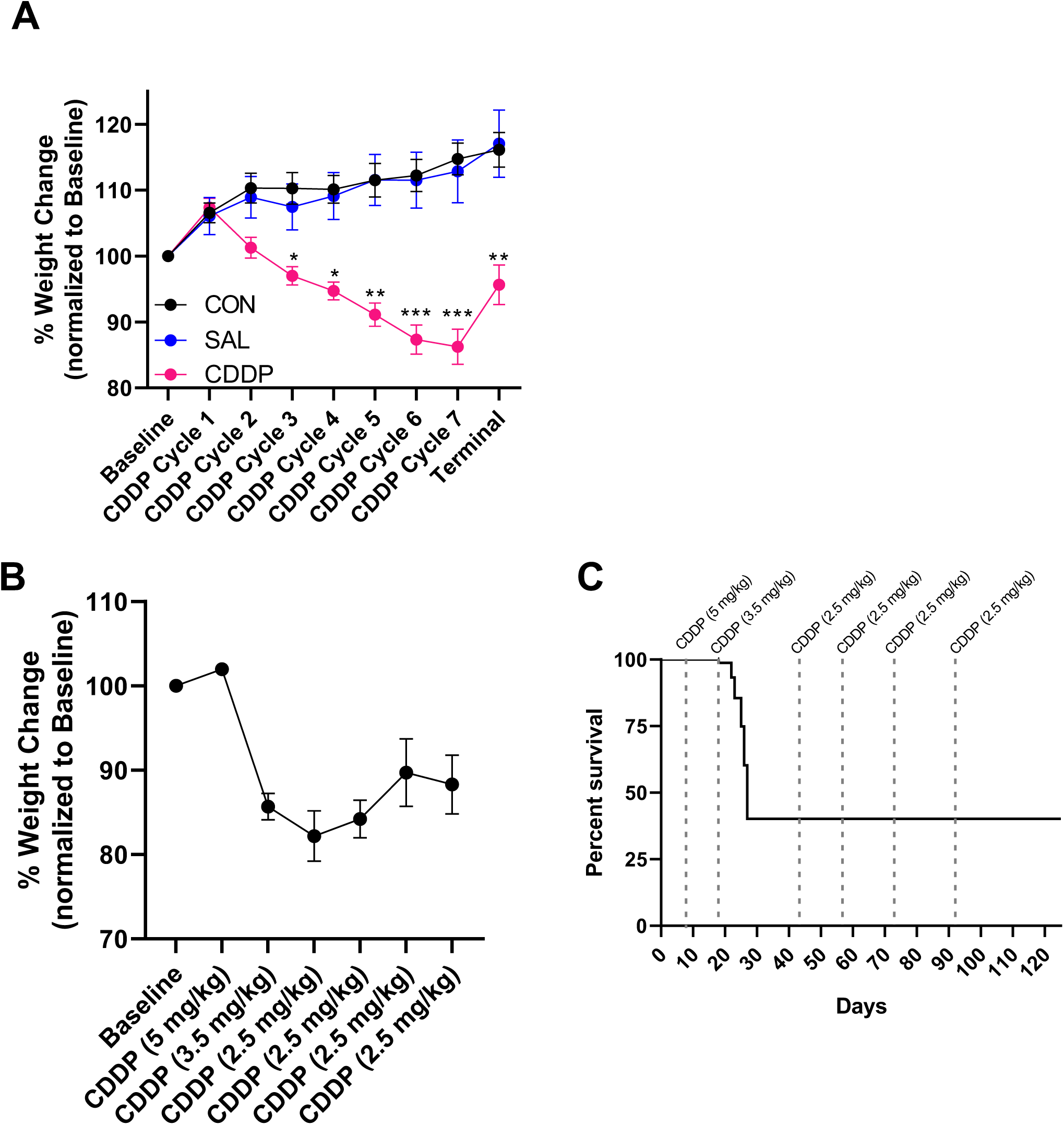
(**A**) Effect of chronic CDDP regimen (2.5 mg/kg, i.p.) every two weeks on body weight (CON: n=8, SAL: n=8, CDDP: n=12). Reported as % in weight change compared to baseline weight. Cisplatin-induced a significant dose-dependent decrease in weight starting CDDP Cycle 3. This CDDP regimen was tolerated, resulting in only a 13.75% maximum reduction in body weight at CDDP Cycle 7, compared to baseline. (**B**) Pilot study to determine the optimal CDDP regimen in middle-aged 9-month old rats (n=12) showed that CDDP (5 mg/kg, i.p) was too high and resulted in a rapid reduction (14.2%) in body weight after the first cycle, and (**C**) 60% reduction in viability after the third CDDP dose. 2.5 mg/kg was determined to be the maximum tolerated dose that could be safely administered every two weeks, without major adverse toxicities.

**Supplementary Figure 2.**
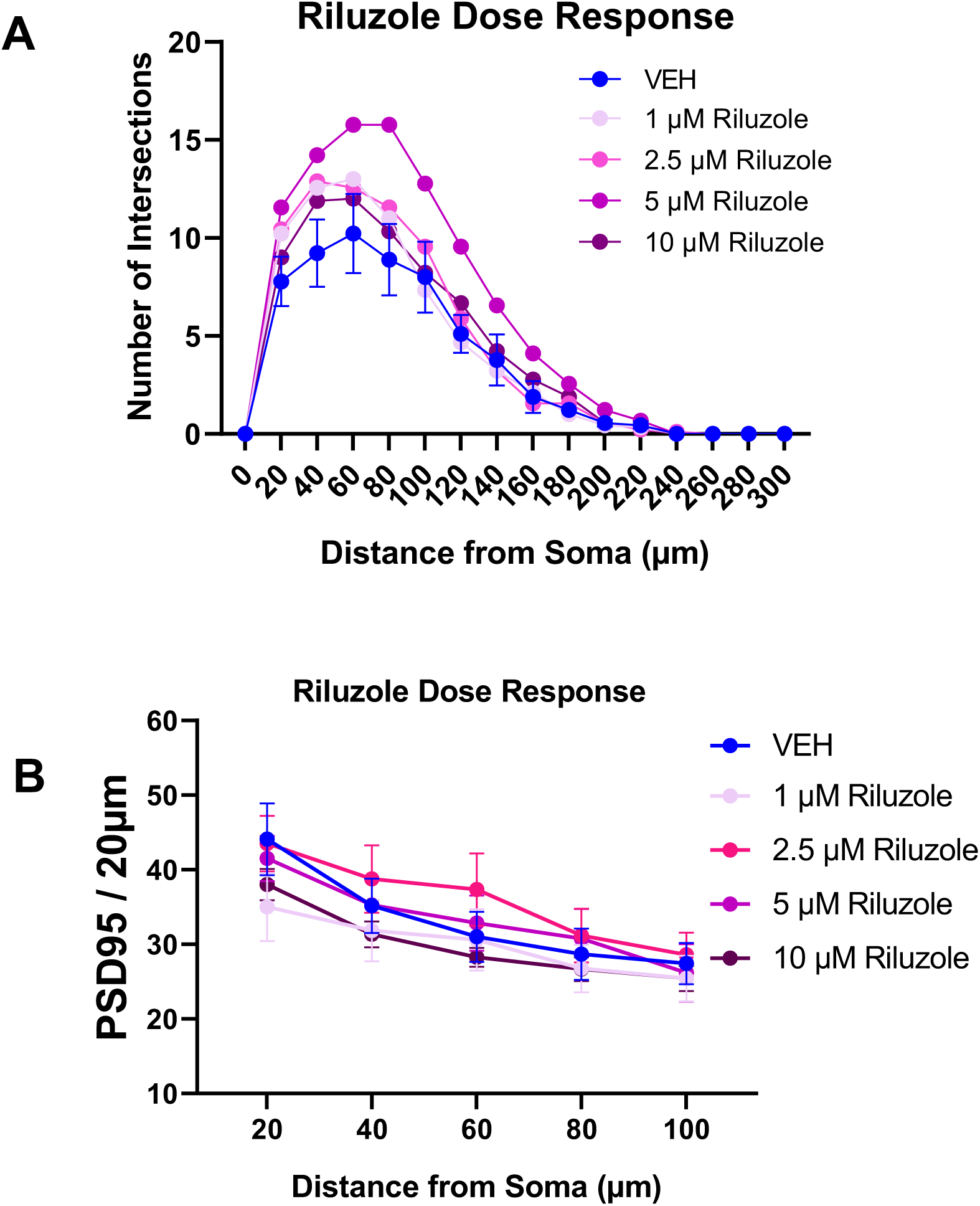
Riluzole dose-response in primary rat hippocampal neurons. Neurons were exposed to graded doses of Riluzole (1 µM, 2.5 µM, 5 µM, 10 µM) for 24 h. (**A**) Quantification of dendritic arborizations of neurons by Sholl analysis. n=9 neurons/group. (**B**) Quantification of PSD95 puncta along 20 µm segments on dendritic branches. n=6 neurons/group, 2 dendritic branches/neuron. Riluzole treatment alone did not significantly change the number of dendritic intersections, two-way RM-ANOVA *F*(_4,40)_ = 1.854, *P*=0.1375, or the PSD95 puncta counts two-way RM-ANOVA: *_F_*(_9,50)_ = 1.693, *P*=0.1156. Data are shown as mean ± SEM.

